# Pink Cedar (*Acrocarpus fraxinifolius*): its prophylactic role against APAP–induced organs toxicity in rats and its antiviral activity against Herpes simplex virus type 1

**DOI:** 10.1101/432328

**Authors:** Eman A. Abdelghffar, Alaa Barakat, Zenab A. Torky, Ihab K. Mohamed, Kamela Ali

## Abstract

The possible protective effects of methanolic extract *Acrocarpus fraxinifolius* leaves (MEAFL) were assessed against the APAP–induced organ toxicity in male rats. Also, the content of polyphenols extracted from AFL was studied, and their relationship with antioxidant activity was investigated. MEAFL was tested for cytotoxicity on Vero cell line, with reference to IC_50_, and other non-toxic concentrations of all the extracts. The antiviral activity against HSV1 for all non-toxic concentrations of the extract was determined using plaque reduction assay. It was found that MEAFL showed a reduction of serum hepatic and renal cellular toxicity and cellular lipid peroxidation, as well as enhanced cellular antioxidant. Also, our results revealed that the inhibitory activity of the virus was dose dependent on the polyphenol content of the examined extract. The MIC for the MEAFL extract was determined as well as the EC_50_ and SI. Calculated SI showed promising value for the MEAFL, and hence can be used as therapeutic medication for HSV1. To study other possible mode of action, Vero cells were treated with the examined extracts before, during, and after virus infection to give an insight on the interference of the extract in each step in the virus life cycle. In conclusion, MEAFL showed a remarkable antioxidant effect against APAP induced organs toxicity. Also, examined extracts exhibited the antiviral activity against HSV1.

## Introduction

Nonsteroidal anti–inflammatory drugs are a non-selective/selective inhibitors for cyclooxygenase (at least two isoforms: COX1 and COX2). Commonly prescribed for their Nonsteroidal anti–inflammatory drugs (such as: aspirin, paracetamol, ibuprofen, naproxen, and diclofenac) are extensively used for the relief of pain, fever and treatment of inflammatory conditions [1,2]. The common mechanism of these drugs is the ability to decrease prostaglandin synthesis by inhibiting COX (COX also known as prostaglandin-endoperoxide synthase). Nonsteroidal anti–inflammatory drugs vary in their relative inhibitory effects on COX-1 and COX-2. There are worried that the COX-2 inhibitors may increase cardiovascular diseases [3]. The overdose of Nonsteroidal anti–inflammatory drugs is associated with an increased toxicity. Also, this toxicity can lead to multiple complications, and may be result in death [4].

N-acetyl-p-aminophenol (APAP), also known as paracetamol or acetaminophen, is a nonprescription drug used as an analgesic, and anti–pyretic drug globally, considered safe and effective at therapeutic doses [2]. APAP toxicity is one of the main causes of poisoning world-wide. Several studies reported that excessive use or overdose of APAP can damage several organs (especially the liver: represents the site of formation of the toxic metabolites, and the kidney: represents the site of its clearance) and even death [2–6] Its toxicity is mediated by the activity of its reactive metabolite (N-acetyl-p-benzoquinoneimine, NAPQI), that generated via cytochrome P450 in liver. NAPQI is detoxified by the antioxidant effects of intracellular glutathione (GSH). Thus, an overdose of APAP cause depletion of cellular GSH [2]. Therefore, it led to a reduced GSH capacity to detoxify NAPQI. Elevation of NAPQI mediates oxidative damage. This subsequently enhances cellular injuries and organ dysfunction [6]. Other studies reported that acute renal damage/failure can occur by overdose APAP even in the absence of liver damage/failure [7,8].

Several studies reported that APAP exerts acute and/or chronic hepato–toxicity, nephro–toxicity, cardiotoxicity effects, gastrointestinal complications and hyperplasia of splenic tissue [1,2,8–13]. 75% of blood advent to the liver arrives directly from gastro-intestinal organs, and then goes to the spleen by portal veins that bring drugs as foreign substances and xenobiotics in near-undiluted form. Following tissue injury, splenic monocytes enter the circulation migrating to inflammatory sites. These splenic monocytes differentiate into macrophages, that participate in pro/anti–inflammatory responses [12]. In dogs, the toxic effects of APAP include hepatic damage, kidney failure and serious hematologic disorders as Heinz bodies formation and hemoglobin damage (non-functioning hemoglobin) [10,14].

On the other hand, HSV-1 is a DNA virus that causes fever blisters, and the primary symptoms of this virus infection are flu-like with fever, followed by the itching and finally those painful papules [15]. Now the real problem with this virus is not about those painful papules, it is about how this virus remains in a latent state in the sensory neurons for a recurrent infection. This recurrent infection, or virus reactivation, is usually triggered by stresses like radiation, and other related factors such as sunlight, menstruation and therapeutic irradiation [16,17]. Hence, if the HSV-1 reactivation is triggered by stress which triggers the oxidative stress, then maybe using antioxidants can cause the oxidative stress to return to the balance state and inhibit the HSV-1 infection. The real problem with herpes is that there is no real cure for it, once a person has the virus, it remains in the body. The virus lies inactive in the nerve cells until something triggers it to become active again. Treatments however can relieve the symptoms and decrease the pain. Treatments can also shorten healing time and decrease the total number of re-infection that is why antiviral agents from plants with new effective compounds exhibiting different modes of action against viral infections are urgently needed.

*Acrocarpus fraxinifolius* leaves (AFL), Fabaceae family and Caesalpiniaceae subfamily, is a native wide spread tree worldwide especially in Africa and Asia. Also, it is distributed in the tropical countries including Egypt. It common name pink cedar, mundani or shingle tree [2,18]. The extracts of the plant were reported to have anti–oxidant, anti–diabetic, anti–proliferative, anti–inflammatory, and hepato– protective activities *in vivo* [2,19–23] as well as antitumor activity *in vitro* [18] [. In general, Caesalpinieae (9 subtribes that have more than 47 genera) have a spectrum of biological activities including anti-oxidant, anti–inflammatory, anti–tumor, anti–diabetic, anti–fungal, anti–bacterial, hepato–protective, gastro–protective, analgesic, anti–arthritic, anti–filarial, anti–malarial, anthelmintic, amoebicidal, diuretic, anti–psoriatic, anti–estrogenic, anti–fertility, wound–healing, anxiolytic, cardio–protective, immune-modulatory,and anti–HIV [2,18–21].

The real value of phenolic compounds is that they possess antioxidant activity which allows them to scavenge free readicals, to inhibit nitrosation, to have the potential for autoxidation and the capability to modulate certain cellular enzyme activities [24]. Acrocarpus fraxinifolius extracts have been proved to possess antioxidant activity [19]. There is little known information about phenolic compounds in MEAFL, their antioxidant properties, and their relation to antiviral activity against HSV-1. Also, there is not enough scanty information on the protective effect of MEAFL on oxidative stress induced by APAP in some tissues like kidney, spleen and heart. So, the objective of this work is to investigate the possible protective activity of MEAFL against APAP–induced organs toxicity (especially kidney, spleen and heart) in male rats, and to examine polyphenolic compounds from MEAFL and then examine the effect of the extract on the cell viability, and conduct the anti-HSV-1 virus activity screening afterwards to check the effect of the extract on virus infectivity. Afterwards, safe concentrations will be selected to determine their effect on each step of the virus life cycle.

## Materials and Methods

### Chemicals

APAP was purchased from Sanofi-Aventisegypts.A.E. (El Sawah, El Amiriya, Cairo, Egypt) and was dissolved in physiological saline containing a minimum amount of dimethylsulfoxide (DMSO, Sigma Chemical Co.). The kits used for biochemical measurements were all purchased from Bio-diagnostic Company (Dokki, Giza, Egypt).

### Preparation of MEAFL

AFL was collected from public garden in Egypt. AFL were prepared by following the previous method [2,23]. 2 kg of AFL was soaked in methanol (80%) for four days then filtered and evaporated till complete dryness.

### Animals

Male Wistar albino rats were purchased from the Animal Breeding House. The animals were housed and fed with pellet diet and tap water ad libitum.

### Experimental design and treatment schedule

Rats were divided into six groups. Group I (healthy control group): rats received orally a 0.2% tween 80 as a vehicle for 21 days; Group II (MEAFL 250): rats received low dose of MEAFL only for 21 days (data not shown). Group III (MEAFL 500): rats received MEAFL only (500 mg/kg b.wt, p.o) for 21 days [2,23]. Group IV (APAP+ vehicle): rats received orally aAPAP (750 mg/kg b.wt; p.o) in the last 7 days to induced oxidative stress in organs [8]; Group V (MEAFL 250 + APAP): rats received orally MEAFL for 21 days then treated with APAP in the last 7 days (data not shown). Group VI (MEAFL 500 + APAP): rats received orally MEAFL for 21 days then treated with APAP in the last 7 days.

### Blood and tissues sampling

Animals were sacrificed after the last administration dose after an overnight fast (on day 22). The blood was collected into tubes with EDTA (for complete blood picture analysis) or without EDTA (for serum markers of cellular toxicity). The kidney, spleen and heart were separated out of the body, cleaned, and weighed then homogenized in 5mL cold buffer (0.5 g of Na_2_HPO_4_ and 0.7 g of NaH_2_PO_4_ per 500mL deionized water, pH 7.4) per gram tissue. Then, the homogenates were centrifuged (15 min/4000 rpm/4 C); and the obtained supernatants were divided into aliquots and preserved at –80°C until used for evaluating the oxidant/anti–oxidant parameters.

### Measurements

#### Evaluation of total phenolic content in vitro

The total phenolic content was determined spectrophotometrically using the Folin–Ciocalteu method based on Slinkard and Singleton, [25], and the early work of Singleton and Rossi, [26].

#### Evaluation of antioxidant activity in vitro

Antioxidant activity was evaluated based on the method reported by Taga et al. [27]. The degradation rate of the extracts was calculated by first order kinetics: Sample degradation rate = ln (a/b) × 1/t.

#### Evaluation of the body weight and organs relative weights of experimental animals

The body/organs weights of liver, kidney & spleen were measured by using a balance (Sartorius LP2200S balance, Gottingen, Germany). Relative weight of organs (g/100 g b.w.) was calculated.

#### Evaluation of the some hematological parameters of experimental animals

Red blood cells (RBCs), hemoglobin (Hb), hematocrit (HCT), white blood cells (WBC), differential granulocytes count (neutrophils, eosinophils and basophils) and differential agranulocytes counts (lymphocytes and monocytes) were determined by automated hematology analyzer (Hemat 8 analyzer; SEAC, Freiburg, Germany). However, blood indices [Mean corpuscular volume (MCV), mean corpuscular hemoglobin (MCH) and mean corpuscular hemoglobin concentration (MCHC)] were calculated from RBCs count, Hb and HCT.

#### Evaluation of serum liver markers of experimental animals

Aminotransferase (ALAT) and aspartate aminotransferase (ASAT) activities were measured according to the method of Reitman & Frankel [28].

#### Evaluation of serum kidney markers of experimental animals

Serum urea was measured according to Vassault et al. [29]. Uric acid was estimated according to Young et al. [30]. Creatinine was estimated by Fossati et al. [31].

#### Evaluation of tissues lipid peroxidation & non-enzymatic/ enzymatic antioxidant status of experimental animals

Thiobarbituric acid reactive substance (TBARS) was assayed according to Ohkawa et al. [32]. The reduced glutathione (GSH) levels were determined according to Beutler et al. [33]. Glutathione peroxidase (GPx) activity was estimated according to the method of Paglia and Valentine [34]. Glutathione reductase (GR) activity was determined according to the method of Goldberg & Spooner [35]. Superoxide dismutase (SOD) activity was estimated according to Nishikimi et al. [36]. Catalase (CAT) activity was assayed by the method of Aebi [37].

#### Cells and Viruses

For the antiviral activity screening, African Green Monkey kidney cells (Vero), were grown in minimum essential medium. Cell cultures were maintained at 37°C in a humidified 5% CO_2_ atmosphere. The virus was propagated in Vero cells, while stock viruses were prepared as previously described Simões et al. [38].

#### Effect of MEAFL on cell viability (cytotoxicity)

The cytotoxic effect of the examined extract on proliferating cells was assayed using an MTT-based method. Toxicity was determined by observation of the morphology of the cells in comparison with the cell control without the extract. After five days of incubation, neutral crystal violet was added for confirmation of cell viability. The cell monolayer was examined by microscopic assessment of changes cell morphology or visible toxic effect [39]. The cells grown in the absence of extracts were used as 100% cell survival. The concentration at which the cell number was reduced to 50% is cytotoxic concentration CC_50_(or inhibitory concentration IC_50_). The concentration which had no or little effect on the cell number, maximum tolerated concentration, or minimum inhibitory concentration (MIC) was also observed.

#### Evaluation of viral plaque number reduction assay

Confluent cell monolayers in 24-well tissue culture plates were adsorbed with 100-200 PFU of HSV for 1h at room temperature. Then, the infected cells were incubated with the different minimum inhibitory concentration of the examined extracts, 230μg/ml for water roots, 86.37μg/ml for water leaves, 72.6μg/ml for acetone roots, and 48.4μg/ml for acetone leaves. Infected cells were then overlaid with medium, containing 1.5% carboxymethylcellulose. After incubation for 5 days at 37°C in 5% CO_2_, infected cells were stained with 0.1% crystal violet, in 1% ethanol, for 15 min. Percentage of viral inhibition after treatment with the extracts was calculated as percentage inhibition compared with untreated viral infected cells control from triplicate experiments.

#### Evaluation of virucidal activity

It was tested by incubation of virus with MEAFL, directly before inoculation of virus onto cells. Briefly, 6-well plate was cultivated with Vero cells, and virus was added to equal volume (V/V) of extract. After 1hr of incubation, the mixture was diluted by 10 fold dilution in a way that still gave suitable count of viral particles. Then the mixture were added to the cell monolayer, and incubated and overlay medium was added. The plates were left to solidify and incubated and observed until the development of the viral plaques. Cell were fixed and stained. Viral plaques were counted and the percentage of viral reduction was calculated [40].

#### Effect of MEAFL extract on Vero cells before HSV-1 adsorption (effect on pretreated cells)

It was tested by subjecting the extract to VERO cells for 2hr before virus inoculation. Briefly, 6-well plate was cultivated with Vero cells and non-cytotoxic concentrations of all examined extracts were added to the monolayer sheet cells and incubated. Extract was removed. Virus was diluted then the virus stock was applied to the monolayer sheet, and then incubated. Overlay medium was added. VERO cells were fixed and stained. Virus inoculums inoculated only to cells and treated identically without addition of the examined extract and served as control. Viral plaques were counted and the percentage of viral reduction was calculated [41].

#### Effect of MEAFL extract on HSV-1 during viral adsorption (effect on attachment and penetration)

Confluent cell monolayers, cultivated in 24-well plates, were infected with virus. Various non-toxic concentrations of the examined extract were added into the cell monolayers and incubated. After that, the inoculum was removed and infected cells were overlaid and incubated. The virus plaques were stained. The EC_50_ was calculated.

#### Effect of MEAFL extract on virus replication

It was tested by post inoculation of extracts after virus application to cells. A 6-well plate was cultivated with Vero cells and incubated. Virus was diluted then the virus stock was applied to the monolayer confluent sheet of cells, and then incubated. Non cytotoxic concentrations with high antiviral activity were applied at different time intervals after virus adsorption. The plates were left to solidify and incubated and observed until the development of the viral plaques. Cell sheets were fixed and stained. Viral plaques were counted and the percentage of viral reduction was calculated [42].

### Statistical analysis

The results were expressed as mean values with their standard errors (±SEM). Data was analyzed using one-way ANOVA, and the differences between groups were determined by Tukey’s multiple comparison test using Graph Pad Prism. Differences at *P*<0.05 were considered statistically significant.

## Results

### Total polyphenolic content and antioxidant activity of MEAFL

Results showed a total of 392.4mg/100g of polyphenol content in the MEAFL, with 85.3% of antioxidant activity.

### Effect of MEAFL on body weight, and organs relative weight of experimental animals

As shown in Fig 1, the body weight gain was significantly decreased (*p*< 0.05) in APAP-intoxicated group (−36.74%) in contrast to the control group. One the other hand, kidney weight and its relative weight were significantly increased (*p*<0.001) in APAP-intoxicated group (61.87% and 72.95%) in contrast to the control group. Oral treatment of MEAFL completely modulated the decrease in in the body weight loss (*p*>0.05) by −22.23% compared to the healthy control animals. Moreover, itsignificantly increased (*p*<0.05) kidney weight (12.83%) and its relative weight (17.01%) compared to the healthycontrol animals. On the other hand, oral treatment of MEAFLsignificantly decreased (*p*<0.01-0.001) kidney weight and its relative weight compared to the APAP-intoxicated group. There were no significant changes in weights of spleen & heart and their relative weights (*p*>0.05) in all treated groups compared to the control animals.

**Fig 1.**
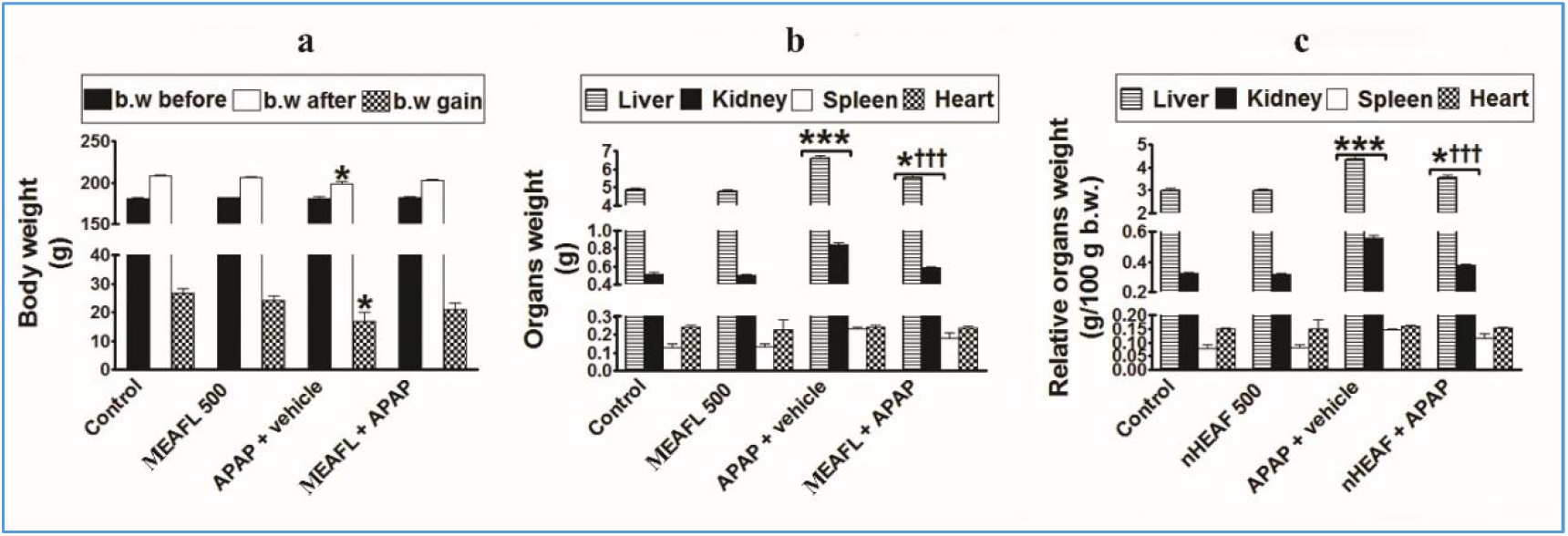
Body weight (a) and organs weight (b) and relative organs weight (c) of control, MEAFL 500 alone, APAP-only and MEAFL500 plus APAP groups.

### Effect of MEAFL on some hematological parameters of experimental animals

There was a slight but not significant decrease(*p*>0.05) in RBCs count, Hb content, and HCT level in APAP-intoxicated group compared with control rats (Fig 2). In addition, granulocytes and agranulocytes were a slight but not significant increase(*p*>0.05) in APAP-intoxicated group compared with the control rats. All these hematological parameters did not significantly change (*p*>0.05) in all treated groups compared to the control animals (Fig 2).

**Fig 2.**
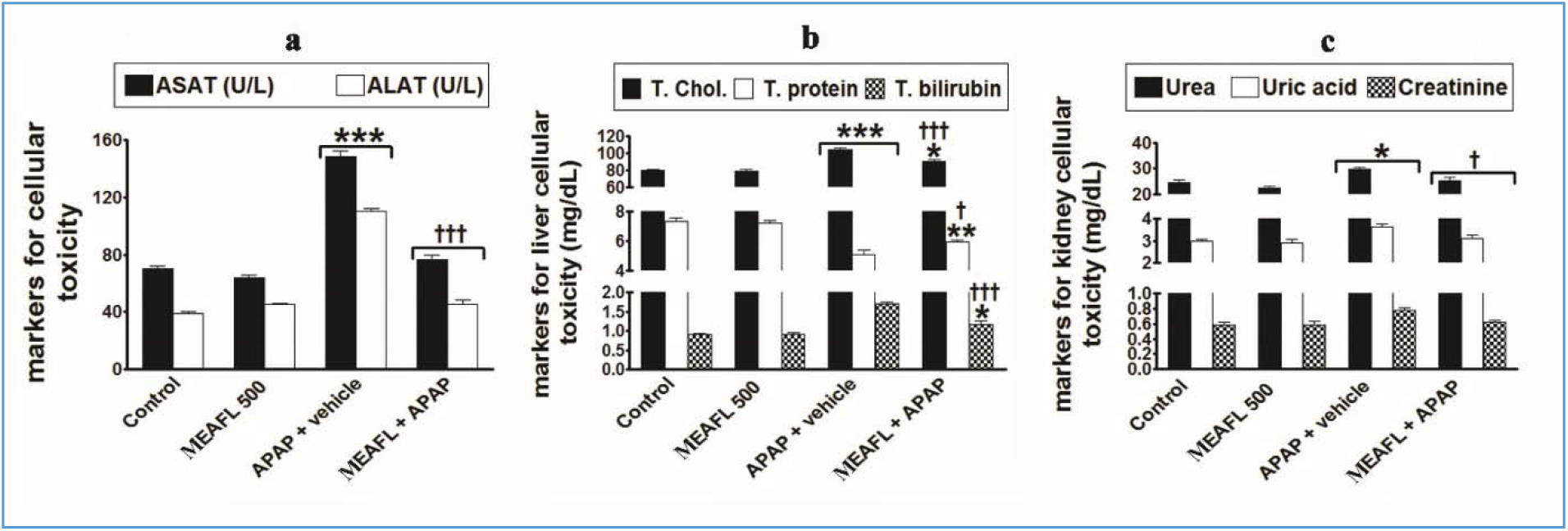
Hematological parameters (a) and blood indices (b), differential granulocytes (c), differential agranulocytes, (d) and total agranulo-agranulocytes counts (e) of control, MEAFL 500 alone, APAP-only and MEAFL500 plus APAP groups.

### Effect of MEAFL on serum hepatic and renal markers of experimental animals

Fig 3 revealed that the serum markers for cellular toxicity (serum ASAT & ALAT activities, urea, uric acid and creatinine levels) were significantly increased (*p*<0.05 to *p*<0.001) in APAP-intoxicated group (112.03%, 184.64%, 21.97%, 25.33%, and 36.61%, respectively) compared with the control animals. Also, oral treatment of MEAFL significanlty increased (*p*<0.05) serum ASAT (9.52%), ALAT (17.65%), urea (3.82%), uric acid (4.18%) and creatinine (4.78%) levels compared with the control animals. Furthermore, oral treatment of MEAFL significanlty decreased (*p*<0.05-0.001) serum markers for cellular toxicity (serum ASAT & ALAT activities, urea, uric acid, and creatinine levels) compared with the APAP-intoxicated control group.

**Fig 3.**
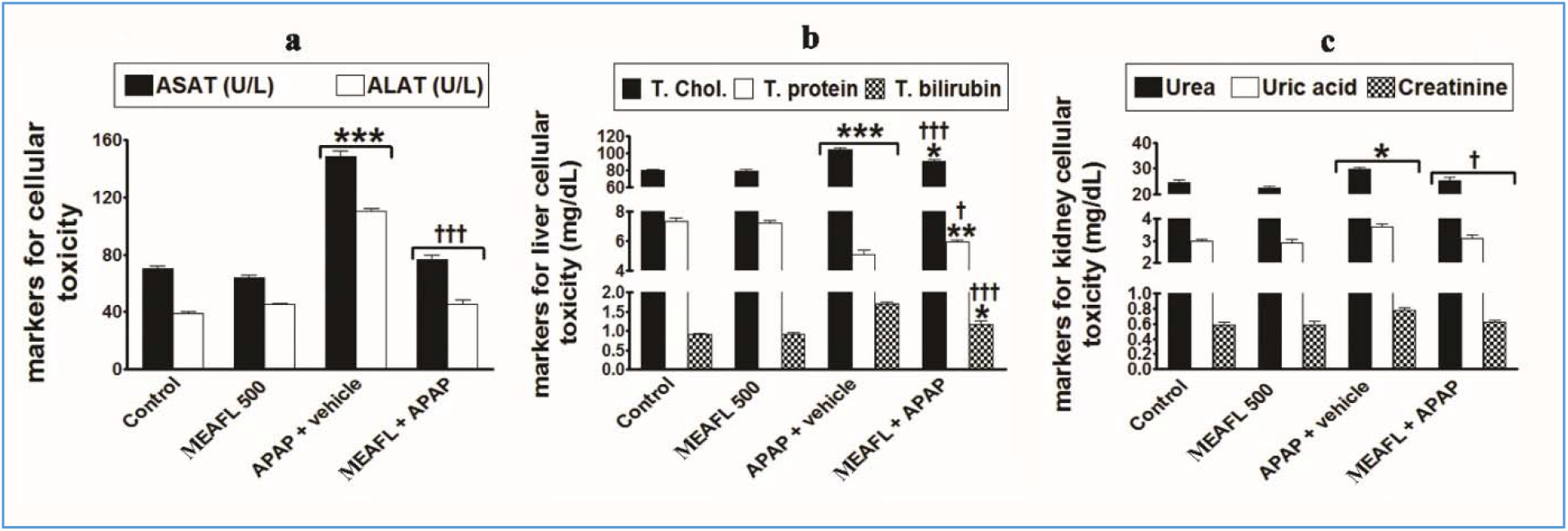
Serum markers for cellular toxicity (a&b) of control, MEAFL 500 alone, APAP-only and MEAFL500 plus APAP groups.

### Effect of MEAFL on cellular oxidant and anti–oxidant markers of experimental animals

As shown in Fig 4, the TBARS content in kidney, spleen and heart was significantly increased (*p*<0.05-0.001) in APAP-intoxicated group (73.35%, 46.36%, and 12.58%, respectively) compared with the control group. Otherwise, the GSH content in kidney, spleen and heart was significantly decreased (*p*<0.001) in APAP-intoxicated group (−57.84%, −34.26%, and −36.81%, respectively) compared with the control group.

**Fig 4.**
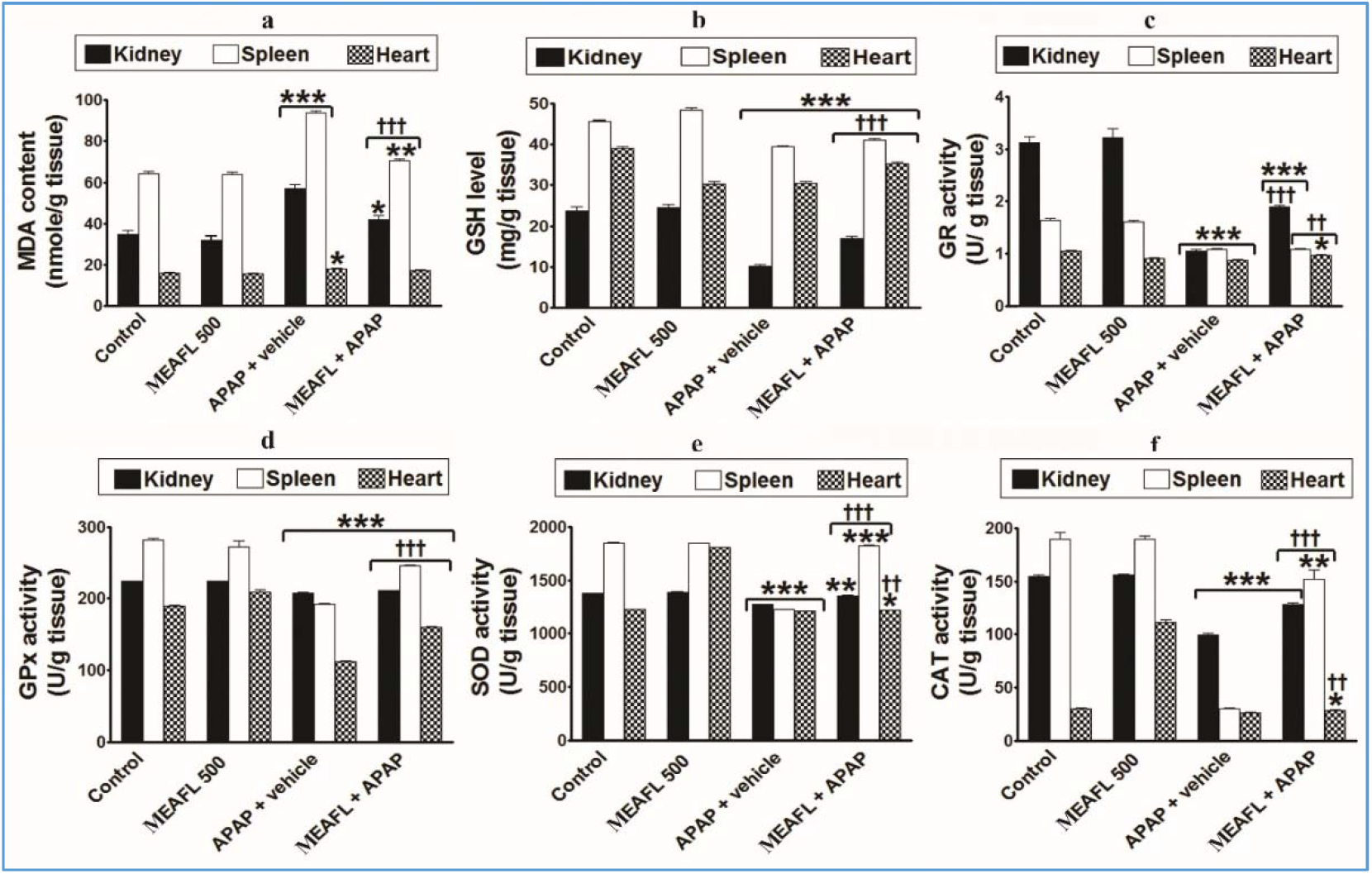
Cellular oxidant and anti–oxidant markers (a-f) in kidney, spleen, and heart of control, MEAFL 500 alone, APAP-only and MEAFL500 plus APAP groups.

Also, enzymatic anti–oxidant (GR, GPx, SOD and CAT) in kidney, spleen and heart were significantly decreased (p<0.001) in APAP-intoxicated group compared with the control group. The percentages of changes of these parameters in tissues (kidney, spleen and heart) of APAP-intoxicated group, compared with the control group, were (GR: −66.93%, −46.42%, &-20.37%), (GPx: −10.16%, −27.90%, &-40.95%), (SOD: −8.14%, −2.33%, &-1.71%), and (CAT: −35.81%, −41.78%, &-24.89%), respectively.

Oral treatmentof MEAFL significantly increased (*p*<0.05-0.01) and decreased (*p*<0.001) TBARS, and GSH, in tissues (kidney, spleen and heart),respectively, but the content of TBARS in heart was not show any significant change (*p*>0.05) in MEAFL plus APAP group compared with the control group. Further, oral treatmentof MEAFL significantly decreased (*p*<0.05-0.001) enzymatic anti–oxidant (GR, GPx, SOD and CAT) in kidney, spleen and heart compared with the control group. Additionally, the percentages of changes of all parameters, in tissues (kidney, spleen and heart)of MEAFL plus APAP group, compared with the control group, were(TBARS:20.54%,10.31%, &7.01%), (GSH:-29.05%,-9.96%, &-19.22%), (GR:-39.39%,-34.05%, &-8.07%), (GPx:-5.75%, −13.01%, &-15.58%), (SOD:-1.60%, −0.92%, &-0.73%), and (CAT:-17.12%, −19.62%, &-11.22%), respectively.

On the other hand, oral treatment of MEAFL significantly decreased (*p*<0.001) TBARS in kidney, & spleen and significantly increased (*p*<0.01-0.001) non-enzymatic and enzymatic antioxidant in tissues (kidney, spleen and heart) compared with the APAP-intoxicated control group.

### Effect of MEAFL on healthy experimental animals

All the above parameters measured in the present study were not significantly altered (*p*>0.05) in healthy-treated rats that received MEAFL compared with the control rats (Figs 1–4). Furthermore, the mortality rates in all groups that received MEAFL were zero during all the period of the study. Subsequently, no deleterious effects were detected for the dose of MEAFL used in this study.

### Effect of MEAFL extract on cell viability (cytotoxicity)

The evaluation of the effects of all examined extracts on the growth and morphology of the Vero cell line using the cytotoxic assay is shown in table (1) and figure (5) below. Two-fold dilutions for AFL lyophilized extract were prepared. All dilutions showed different degrees of the CPE especially on higher concentrations, while the lower concentrations showed no CPE on the Vero cell line with IC_50_ of 108.9 μg/ml and cell-safe concentrations <=72.6 μg/ml.

**Figure 5:**
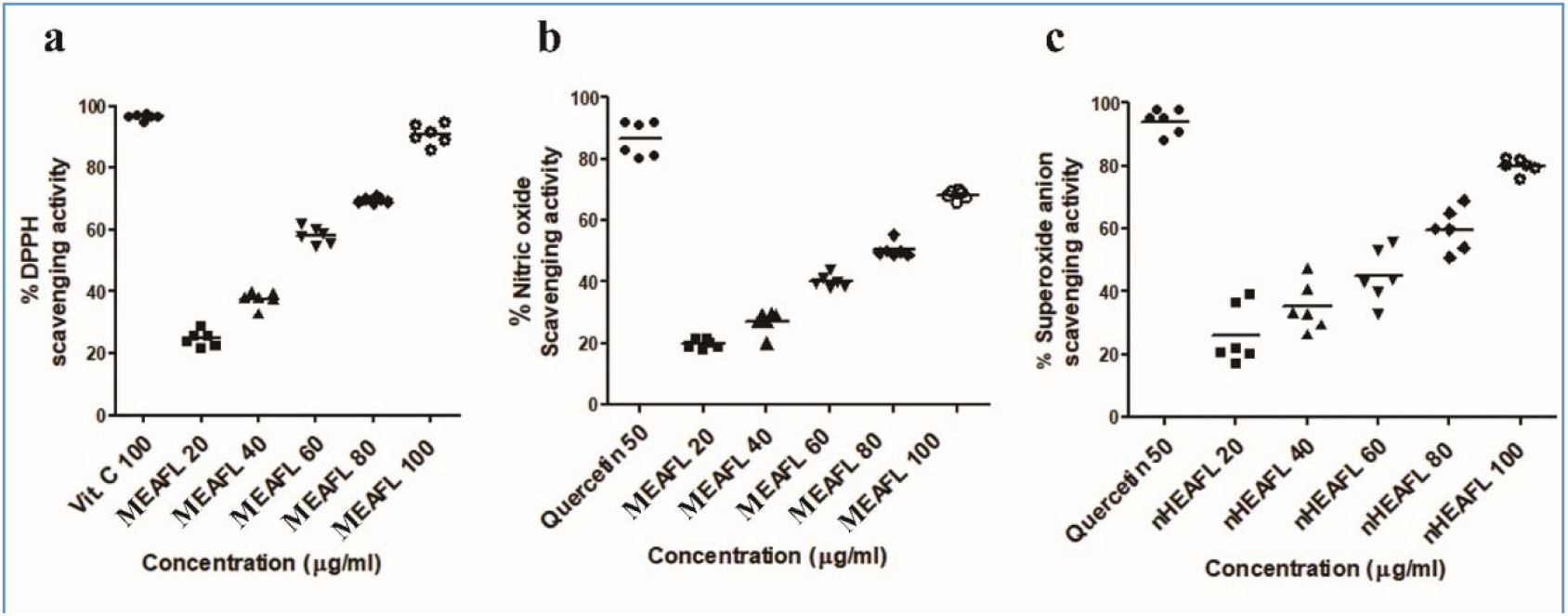
In vitro free radical scavenging effect of MEAFL in vitro by DPPH (a), nitric oxide (b), superoxide radicals scavenging (c) methods.

**Figure 5:**
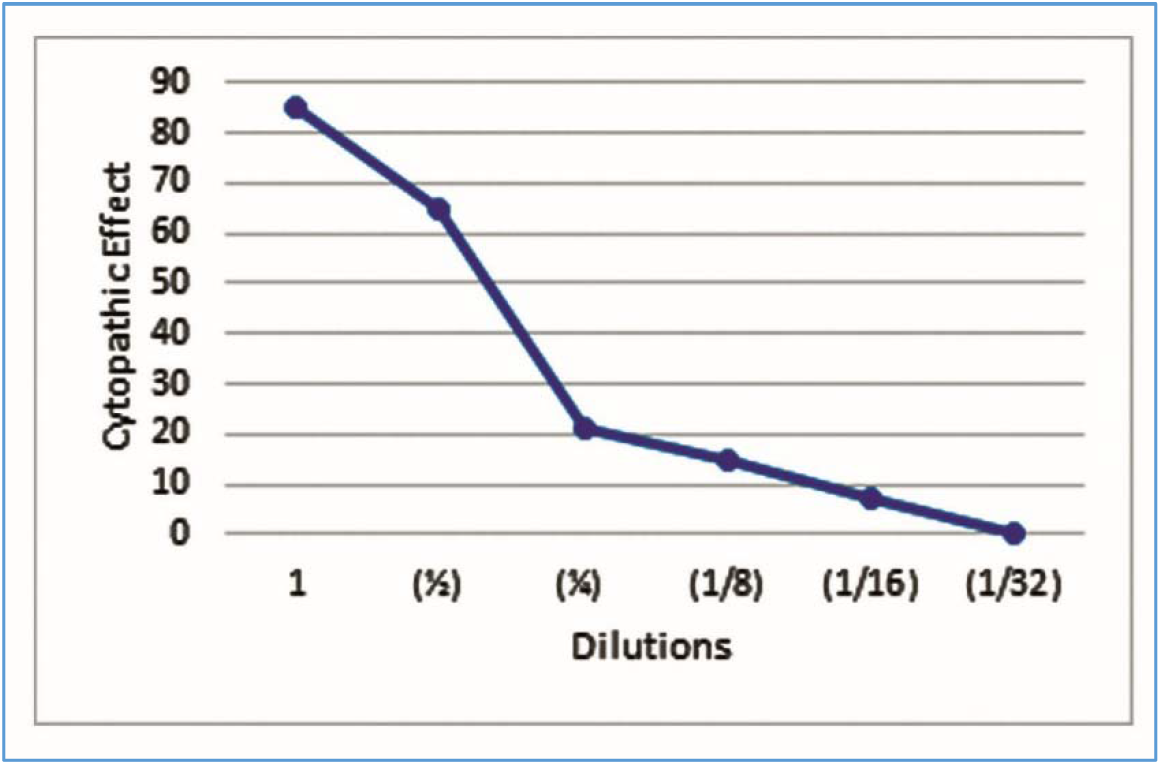
CPE % of MEAFL extract.

**Table 1:** CPE % of MEAFL extract.

| **Dilutions** | 1 | 1/2 | 1/4 | 1/8 | 1/16 | 1/32 |
| --- | --- | --- | --- | --- | --- | --- |
| **CPE %** | 85.3 | 64.5 | 21.3 | 15.1 | 7.5 | 0 |

### Viral plaque number reduction assay

The experiment of the cytotoxic effect of the MEAFL extract on the Vero cell line gave the safest concentrations, and those concentrations were used in the evaluation of the antiviral activity of the extracts in the following experiments to study the effect of the extract on the virus life cycle, and determine the EC_50_, which is the 50% viral inhibitory concentration, and the selectivity index (SI) which is calculated as the IC_50_/EC_50_ and finally, to choose the minimum inhibitory concentration (MIC) that is the concentration with no or little cytotoxicity and maximum antiviral activity.

### The antiviral activity of the MEAFL extract was assayed by plaques number reduction assay

Results in table (2), and figure (6) below show that the extract started with a moderate antiviral activity of 67% at dilution 1/4 which is equivalent to concentration 72.6 μg/ml. This antiviral activity almost disappeared at dilution 1/8 giving 39.2% antiviral activity, the extract has EC_50_ of 51.9 μg/ml, SI of2.09 and MIC of 72.6 μg/ml.

**Figure 6:**
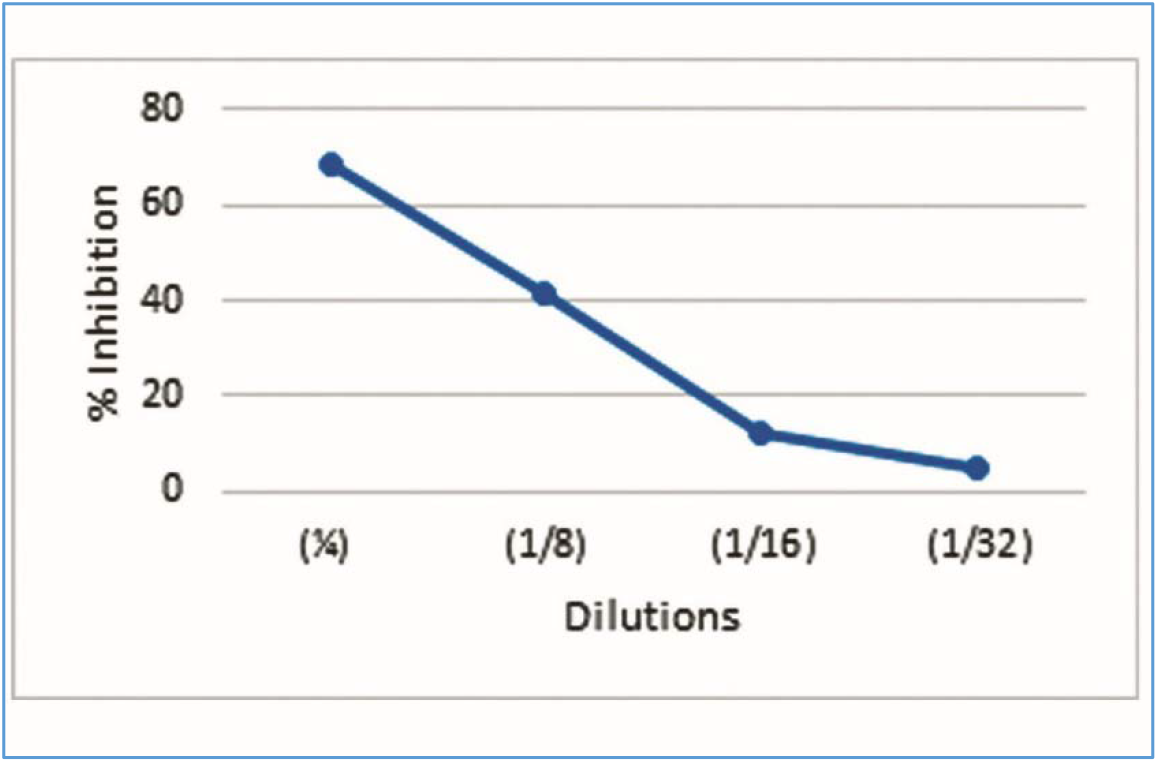
PRA of each cell-safe dilution of MEAFL extract.

**Table 2:** PRA of each of the cell-safe dilutions of MEAFL extract.

| **Dilutions** | 1/4 | 1/8 | 1/16 | 1/32 |
| --- | --- | --- | --- | --- |
| **% Inhibition** | 68.4 | 41.2 | 12.3 | 5.3 |

### Virucidal Activity

After determination of toxic and safe concentrations of MEAFL extract on Vero cell line, MIC concentration was incubated with HSV-1 in cell free culture media for one hour. Results showed a total inhibition of 87% of the HSV-1 virus.

### Effect of MEAFL extract on pretreated cells

To determine the mode of antiviral action, the examined AFL extract, was added to the Vero cells at different time intervals before, during, and after the virus adsorption.

AFL MIC concentration was added to the Vero cells for 1 hour and completely removed by washing the cells, before HSV-1 inoculation. PRA showed that the extract showed a very low degree of virus inhibition around 8.4%.

### Effect of MEAFL extract on HSV-1 during viral adsorption (effect on attachment and penetration)

AFL MIC concentration was added at the same time of the HSV-1 inoculation. Results showed an inhibition of 71.2% of the HSV-1 virus infection.

### Effect of MEAFL extract on virus replication

Effect of MEAFL MIC concentration on viral replication was demonstrated at different time intervals (1, 4, 8, 24 and 48 hours) after virus inoculation. Results are shown in table (3) and figure (7) below.

**Figure 7:**
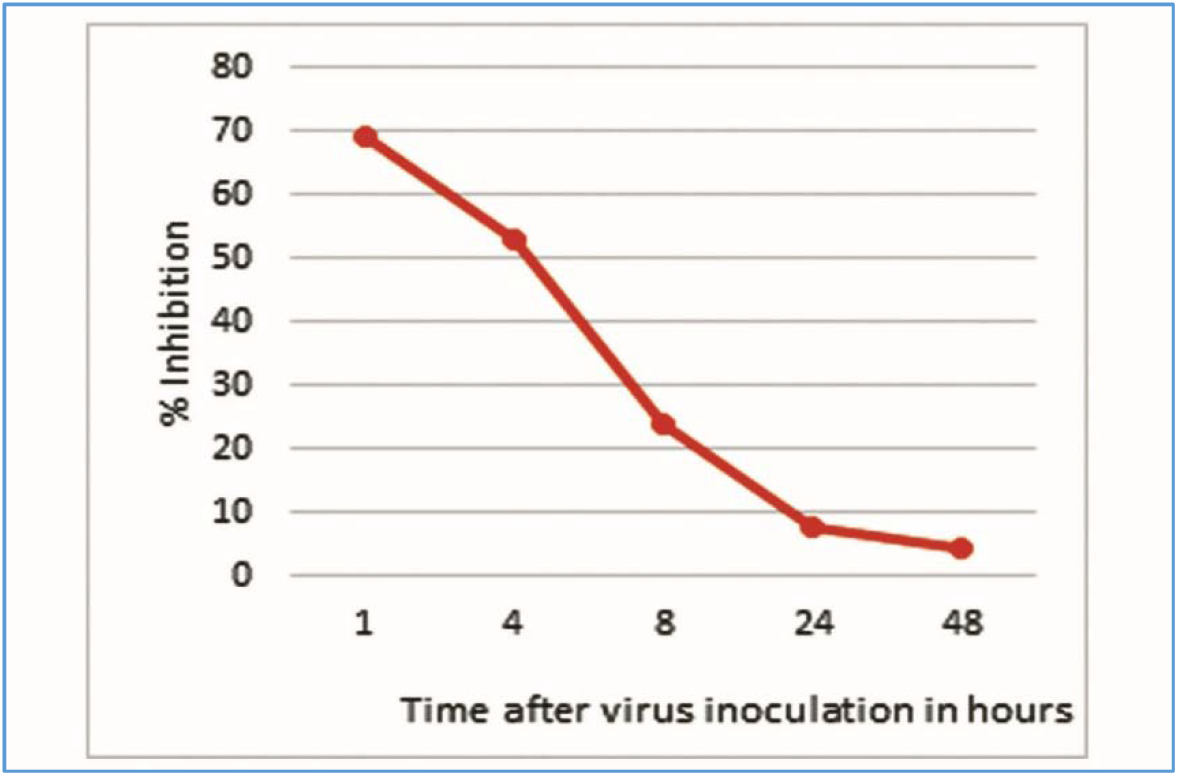
Anti-HSV of MEAFL extract at different time intervals after virus inoculation.

**Table 3:** Anti-HSV of MEAFL extract at different time intervals after virus inoculation.

| **Time after virus inoculation in hours** | 1 | 4 | 8 | 24 | 48 |
| --- | --- | --- | --- | --- | --- |
| **% Inhibition** | 68.5 | 53 | 23.4 | 8.6 | 5.3 |

## Discussion

As for the in vitro experiments, the MEAFL was examined for polyphenol contents and revealed a high content, Some studies encourage the use of different extraction systems like ethanol [43], and methanol [44] to extract polyphenols with antioxidant activities. Some other studies preferred the use of water extraction methods of polyphenols [45–47]. Comparable results were also reported by Trouong et al. [48] on the sweet potatoes. Results also showed that when polyphenols content increase, a noticeable increase in antioxidant activity happens. Same results were reported by Donglin, and Yasunori [49].

In vivo experiments, pretreatment with MEAFL produced partial protection against overdose APAP-induced hepatotoxicity, nephrotoxicity, cardiotoxicity, and splenotoxicity, represented by significant reduction of serum cellular markers,cellular lipid peroxidation (TBARS) and significant elevation cellular anti-oxidants (GSH, GR, GPX, SOD and CAT) compared with non-treated APAP group. Thus, it was determined the ability of MEAFL’s to protect against cellular GSH depletion, restore the pro-oxidative/antioxidative cellular imbalance, and prevent lipid peroxidation [22,23]. In addition, the reversal of increased serum cellular markers in APAP-induced organs injuries by MEAFL may be due to the prevention of the leakage of intracellular enzymes by its membrane-stabilizing activity [2, 22,23]. Also, these modulations led to alleviate the loss in body weight gain [2] and relative kidney weight. This may be attributed to the fact that phenolic acids in AFL have anti–oxidant and free radical chelators/scavenger’sactivities as with special impact over hydroxyl/peroxyl radicals, superoxide anions, and peroxynitrites [18].

Recently, bioactive phenolics such as brevifolin carboxylic acid, ellagic acid, gallic acid and methyl gallate were identified from the extract of AFL [18,20], and may be responsible for its radical scavenging activity [50].

Our previous studies have indicated that MEAFL (500 mg/kg,p.o, for 7 to 21 days) plays an important role in improving GSH status and total anti–oxidant capacity in intoxicated rats with APAP [2,23]. In this study, the high level of GSH in response to MEAFL may result from increased activity of cellular anti–oxidant enzymes and decreased cellular lipid peroxidation.

Higher activity in MEAFL may be due to presence of α–tocopherol which has a powerful anti–oxidant activity in detoxifying free radicals, stabilization of the cell membrane and structure restoration

[2]. α–Tocopherol may have inhibited the chain reactions of APAP–generated free radicals or scavenged the ROS before reaching its renal, splenic and cardiac targets. Furthermore, α–tocopherol stimulated the upregulation of endogenous cytochrome P3 (A4 and A5) which metabolize APAP into reactive metabolite NAPQI [2,51].

The Polyphenols, flavonoids, and anthocyanins are known to possess antioxidant activities (scavenging of free radicals), due to their several phenolic hydroxyl groups [52]. Also, H_2_O_2_ scavenging by polyphenolic extract can be attributed to its phenolic nature, which can give electrons to H_2_O_2_ and thus neutralize it to water [52]. Recently, El-Kashak et al. [20] reported that AFL have very strong anti–oxidant effect due to presence of flavonoids (such as: quercetin–3–O–β–D–glucopyranoside, quercetin–3–Oα–L–rhamnopyranoside, myricetin-3–O–β–D–galactopyranoside and myricetin–3–O–α–L–rhamnopyranoside). Also, the phytochemical screening of MEAFL demonstrated the presence of polyphenolic components; such as α-tocopherol, labda-8 (20)-13-dien-15-oic acid, lupeol, phytol and squalene [2]. Also, it has been reported that MEAFL contains flavonoids, tri–terpenoids, α–tocopherol and steroids, which exhibited strong anti–oxidant activity. Several authors have already confirmed that gallic acid and α-tocopherol are potent antioxidants [2,50,53]. Another scientific report indicated certain flavonoids, tri-terpenoids and steroids have the protective effect on hepatic tissue due to its anti–oxidant properties [54]. The presence of polyphenolic compounds in MEAFL may be responsible for the protective effect against APAP-induced organs toxicity in rats.

Abd El-Ghffar et al. [2] and Alaa [23] proved that the anti–oxidant and hepato–protective activities exerted by MEAFL against APAP-induced oxidative damage was attributed to its active constituents such as lupeol, squalene, and phytol. Also, another study proved that lupeol (from *Ficuspseudopalma*) has the anti–oxidant and hepato–protective activities against APAP-induced oxidative damage [55]. Sivakrishnan & Muthu [56] and Zuhan et al. [57] reported about the promising hepato–protective effects of squalene (isolated from *Albiziaprocera*) against APAP– and CCl_4_–induced toxicity. Also, it has been reported that phytol acts as a precursor for vitamin E and K1 and it has anti–oxidant as well as anticancer activities [2,58]. Taking this fact with gained results we could suggest that the amount of exogenous polyphenolic diets increases, endogenous anti–oxidant defense system increase as well. Therefore, the medicinal anti–oxidant therapeutic will offers a promising way to prevent and/or treat the diseases induced by the excessive exposure to oxidative stress (ROS/NOS).

From our in vivo study, we can reported that MEAFL has important anti–oxidant effects as ability to scavenge ROS, inhibit lipid peroxidation as well as increase of anti–oxidant defense system in kidney, spleen, and heart tissue organs; which was attributed to its active constituents.

On the other hand, before any experiment can be conducted on the MEAFL to study its effect on HSV-1, or its mode of action, the cytotoxicity of the extract on Vero cells has to be studied first. Dilutions of the MEAFL were examined for biological activity toward the viability of Vero cell line. Determination of cytotoxicity of the used extract is an important factor in the evaluation of any antiviral substance. As efficient SI values should have very little CPE on the cells, and very high antiviral effect on the virus, It can be concluded from the results of the viral plaque number reduction assay experiment results that the MEAFL has a very promising results as a therapeutic treatment for HSV-1, since almost all of its concentrations used were safe and consequently a high concentration can be used which will give a high antiviral activity and a strong effect as a therapeutic agent

Results of the virucidal activity of the MEAFL against the HSV-1 virus showed that considerable amount of the virus inhibition was due to direct virus inactivation due to the direct contact with the virus particles. The degree of viral inhibition is directly correlated with the concentration of the polyphenol of the extract. Those results come in accordance with [40,59–61] who stated that the exposure of the cell free HSV-1 to black berry for 15mins in room temperature had a virucidal activity.

As for the mode of action experiments, the fact the MEAFL pretreated Vero cells showed a very low viral inhibition, can lead to a conclusion that pretreatment of cells of tissue culture with the examined MEAFL before virus inoculation has no effect on virus infectivity. Comparable results were reported by Raenu et al. [62], who concluded their research stating that the extracts they used did not have any effect on the HSV-1 virus infectivity when used on the Vero cells before the virus inoculation.

On the other hand, the results of viral inhibition during viral adsorption indicate that the examined extract inhibited the virus replication at different degrees according to the concentration via blockage of the adsorption of the virus either by the deactivation of binding of the surface glycoprotein of the viral envelope of virus particles to the cell receptor sites or inhibiting the viral cell fusion in the early replication stage and finally, preventing the initial stages of the viral reproduction. Same results were reported by previous studies [59, 63, 64], who stated that the extracts they used directly inactivated HSV-1 particles, leading to the failure of early infection, including viral attachment and penetration [65].

When antiviral activity of MEAFL against HSV-1 was measures at different time intervals after virus inoculation, results show that the extract caused different degree of inhibition depending on the time of addition. Also, treatment of Vero cells one to four hours after virus inoculation showed the highest virus inhibition.

From the in vitro results, it can be concluded that, as the time interval increases, between the virus adsorption and addition of the extract, the inhibition of virus activity decreases significantly after four hours until it vanishes. These results indicate that the viral inhibition observed at the early hours just after the virus inoculation happens via the interference of the examined extract with the viral protein biosynthesis and transcription through the early phase of viral replication. Comparable results were reported by Sayed et al. [66].

## Conclusion

The results concluded that oxidative stress is participated in the pathogenesis of APAP-induced organs toxicity. MEAFL possesses marked hepato-, nephro-, spleno- and cardio-protective activities against overdose of APAP induced organs toxicity. The protective effect of AFL may be due to its phytoconstituents that had a powerful antioxidant activity and free radical scavenging properties. Subsequently, it is suggested to use these medicinal plants as a food supplements in diet to alleviate versus toxic effects of APAP. Therefore, further studies will be required to determine the exact mechanisms, by which it exert their effects, associated with our findings. Furthermore, results of the in vitro experiments showed a great therapeutic potential of the MEAFL with a very low to noncytotoxic effect on Vero cells.

## Availability of data and materials

The datasets supporting the conclusions of this article are included within the article and its additional files.

